# Super-Resolution Cryo-EM Maps With 3D Deep Generative Networks

**DOI:** 10.1101/2021.01.12.426430

**Authors:** Sai Raghavendra Maddhuri Venkata Subramaniya, Genki Terashi, Daisuke Kihara

## Abstract

An increasing number of biological macromolecules have been solved with cryo-electron microscopy (cryo-EM). Over the past few years, the resolutions of density maps determined by cryo-EM have largely improved in general. However, there are still many cases where the resolution is not high enough to model molecular structures with standard computational tools. If the resolution obtained is near the empirical border line (3-4 Å), a small improvement of resolution will significantly facilitate structure modeling. Here, we report SuperEM, a novel deep learning-based method that uses a three-dimensional generative adversarial network for generating an improved-resolution EM map from an experimental EM map. SuperEM is designed to work with EM maps in the resolution range of 3 Å to 6 Å and has shown an average resolution improvement of 1.0 Å on a test dataset of 36 experimental maps. The generated super-resolution maps are shown to result in better structure modelling of proteins.

## Introduction

Technological advances in cryo-EM have led to its rapid adoption in solving structures of biological macromolecules, including structures that were challenging to determine by other experimental methods^1^. Particularly, owing to recent software and hardware breakthroughs in this field, an increasing number of structures have been solved at atomic or near-atomic resolutions^2^. However, there are still many cases where maps are determined at a lower resolution where structure modeling^3,4-7^ is not straightforward. Since structure modeling of macromolecules becomes increasingly difficult as map resolution worsens, particularly across the critical resolution range of 3 to 4 Å, an improvement of a resolution even by 0.5 Å to 1 Å is valuable. The improvement of map resolution is commonly attempted experimentally by altering map building steps and data collecting steps, including accumulating more two-dimensional particle images, but this task is time-and labor-intensive.

There are some computational methods developed for post-processing cryo-EM maps through local sharpening approaches. LocScale^8^ makes use of solved atomic models to upscale the map densities. LocalDeblur^9^, is a local sharpening method that models the relation between experimental map and sharpened map. Terwilliger et al. developed a maximum-likelihood approach for density modification approach from half maps^10^. DeepEMhancer used deep learning for sharpening maps^11^.

Here, we propose a novel 3D deep learning based super-resolution method, SuperEM, which uses generative adversarial networks (GAN)^12^ to improve the resolution of experimental EM maps in the resolution range of 3 Å to 6 Å. Super-resolution imaging is a task of estimating high-resolution images from their low-resolution counterparts^13^. This task is traditionally addressed with image interpolation techniques^14^ or by considering image statistics^15,16^ and local image patches^17^. With the advancements in deep learning, the recent introduction of GAN has achieved state-of-the art performance in improving the resolution of images^18^. The same framework has been successfully applied to various interesting tasks, such as improving medical imaging^19,20^, image restoration^21^, image compression^22^, and image-to-image translation^23^. Super-resolution imaging applications of GANs were also extended to other microscopy image data, such as from fluorescence microscopy^24^, single-molecule localization microscopy (SMLM)^25^, and CT imagery^20^.

Our method, SuperEM, uses a 3D GAN to produce super-resolution 3D EM maps from an input low-resolution map. SuperEM GAN consists of two neural networks, generator and discriminator, which are trained simultaneously. SuperEM was trained with a dataset of pairs of an experimental EM map coupled with a corresponding high-resolution 3D simulated EM map that was generated from the associated protein atomic structures. The goal of training was to make the generator of SuperEM be able to output high-resolution EM maps that are not distinguishable from actual high-resolution maps by the discriminator network. At the same time, the discriminator was trained to distinguish between the generated and real high-resolution maps.

SuperEM is shown to produce EM maps with substantially higher quality than the input low-resolution experimental maps. An average resolution improvement of 1.0 Å was observed when tested on a test dataset of 36 experimental EM maps. Furthermore, maps produced by SuperEM also led to better protein structure models when modelled with MAINMAST^3^ and Phenix^5^.

## Results

### GAN architecture of SuperEM

SuperEM receives a low-resolution 3D cryo-EM map as input and produces a super-resolution (SR) map as output. SuperEM adopts a network architecture similar to SRGAN^18^. It was trained using a low-resolution (LR), experimental EM map and a corresponding high-resolution (HR) EM map simulated from the associated atomic-detail structure of proteins. The inputs to GAN are a pair of cubes of a volume size of 25^3^ Å^3^ each, which are extracted by scanning the EM maps. The generator module of the GAN outputs an SR cube of the same size. Individual SR cubes output from the network are combined to generate a complete super-resolution EM map.

**Fig. 1** illustrates the GAN architecture of SuperEM. The generator network is a fully convolutional network consisting of a 3D convolution layer with 32 channels and a kernel size of 3 followed by 15 residual network (ResNet) block layers^26^, a 3D convolution layer ending with a tanh activation layer. Each ResNet block contains two 3D convolutional layers with 32 channels and skip connections, dropout (with a dropout probability 0.25), PReLU activations^27^, and Instance Normalization^28^, which was shown to work better than other normalization methods for generative tasks. The final 3D convolutional layer also uses a kernel size of 3 with 32 output channels. The last tanh layer outputs a probability distribution for each voxel in the cube, which is interpreted as the density distribution in the cube. The output of tanh function is normalized to produce values in the range of 0 to 1. A stride of 1 was used in all the filters in all the layers.

**Figure 1.**
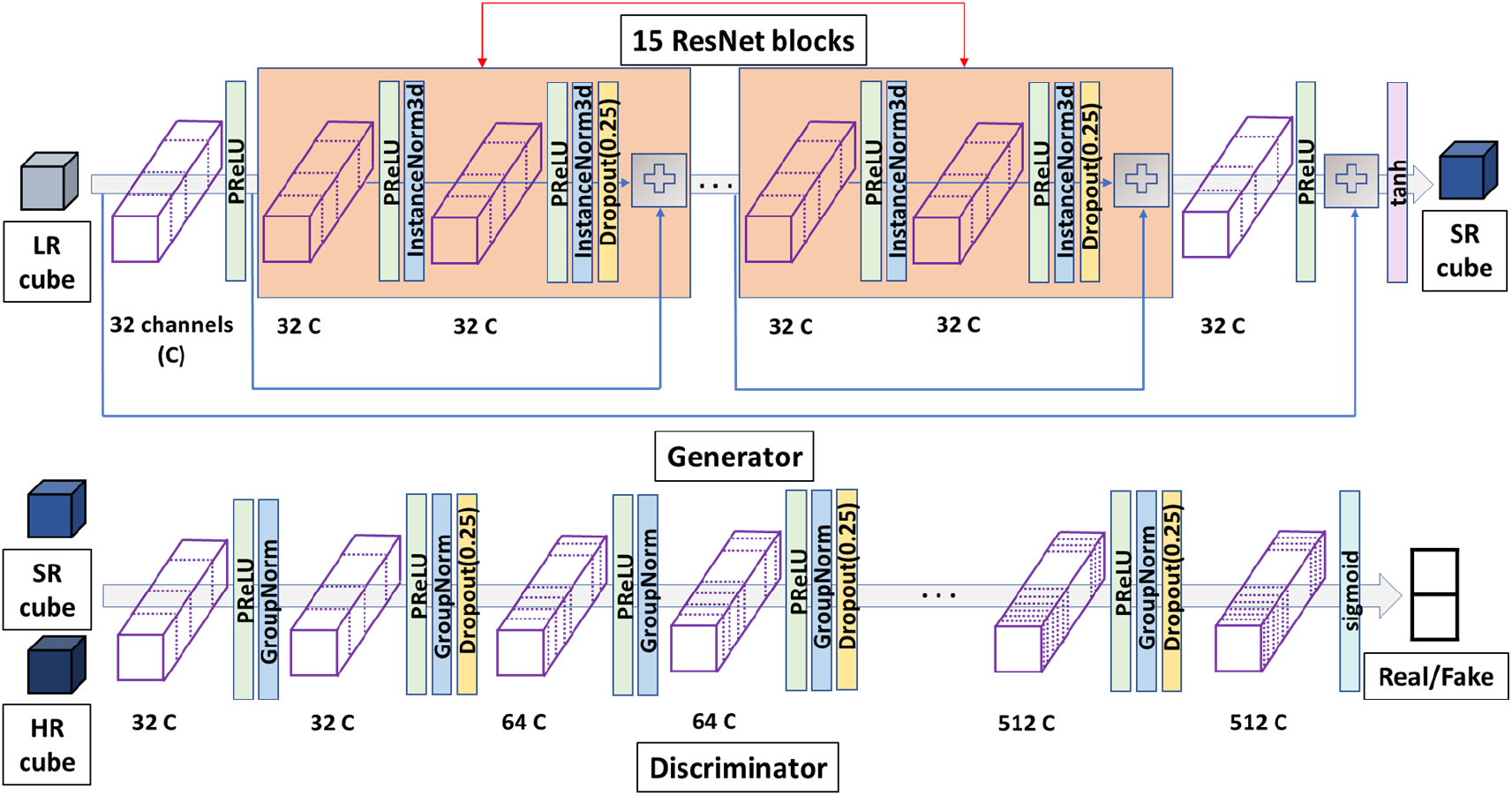
The 3D GAN architecture of SuperEM. The detailed architecture of the generator and discriminator networks are shown. LR, low-resolution; SR, super-resolution; HR, high-resolution; PReLU, the parametric rectified linear unit activation function; InstanceNorm3D, 3D instance normalization layer; GroupNorm, group normalization layer. The blue arrows that connect the input of a ResNet block and a plus sign, which is the operator that simply add two matrices, is a skip connection. See text for more details.

The discriminator network is a fully convolutional binary classifier. It takes a generated map output from the generator and the corresponding HR density map and classifies the two maps into classes, real or generated (fake). The discriminator network consists of 10 3D CNNs followed by a softmax layer. The first convolutional layer in the discriminator contains 32 channels and the count is multiplied by a factor of 2 for every two subsequent layers. We use PReLU, dropout with a probability of 0.25 and group normalization with a group size of 16 in all the convolutional layers.

We evaluated the quality of EM maps generated by SuperEM on a dataset of 36 EM maps from multiple angles. The resolution of the test maps ranged from 3 Å to 6 Å with 17 maps between 3 Å-4 Å and 14 maps between 4-5.5 Å. In terms of protein secondary structure contents, nine maps were helix-rich (over 40% of the total residues are from α helices), nine maps are sheet-rich (β strand residue counts > 40%), and 18 maps with about equal distribution of helix and β strand residues (both α helix and β strand residue counts > 25%). See Methods for detailed descriptions of train/test datasets, training procedure, and the evaluation criteria.

### Improvement of local map regions by SuperEM

We first evaluate the improvement of density maps by SuperEM at the local cube level. Our data generation procedure, as described in Methods, generated 91,197 cubes of size 25^3^ Å from test maps. These low-resolution (LR) cubes are fed to SuperEM GAN, which generated the same number of corresponding super-resolution cubes (SR). As references, we generated high-resolution (HR) cubes taken from the simulated maps that were computed from atomic structures from PDB^29^. For experimental maps of a resolution between 3 Å and 3.5 Å, HR maps were simulated at a 1.8 Å resolution while those between 3.5 Å and 6 Å, a HR map was simulated at 3 Å. We then calculated cross-correlation between LR/HR cubes and SR/HR cubes using the Chimera’s “measure correlation” tool^30^ (**Figure 2a**).

**Figure 2.**
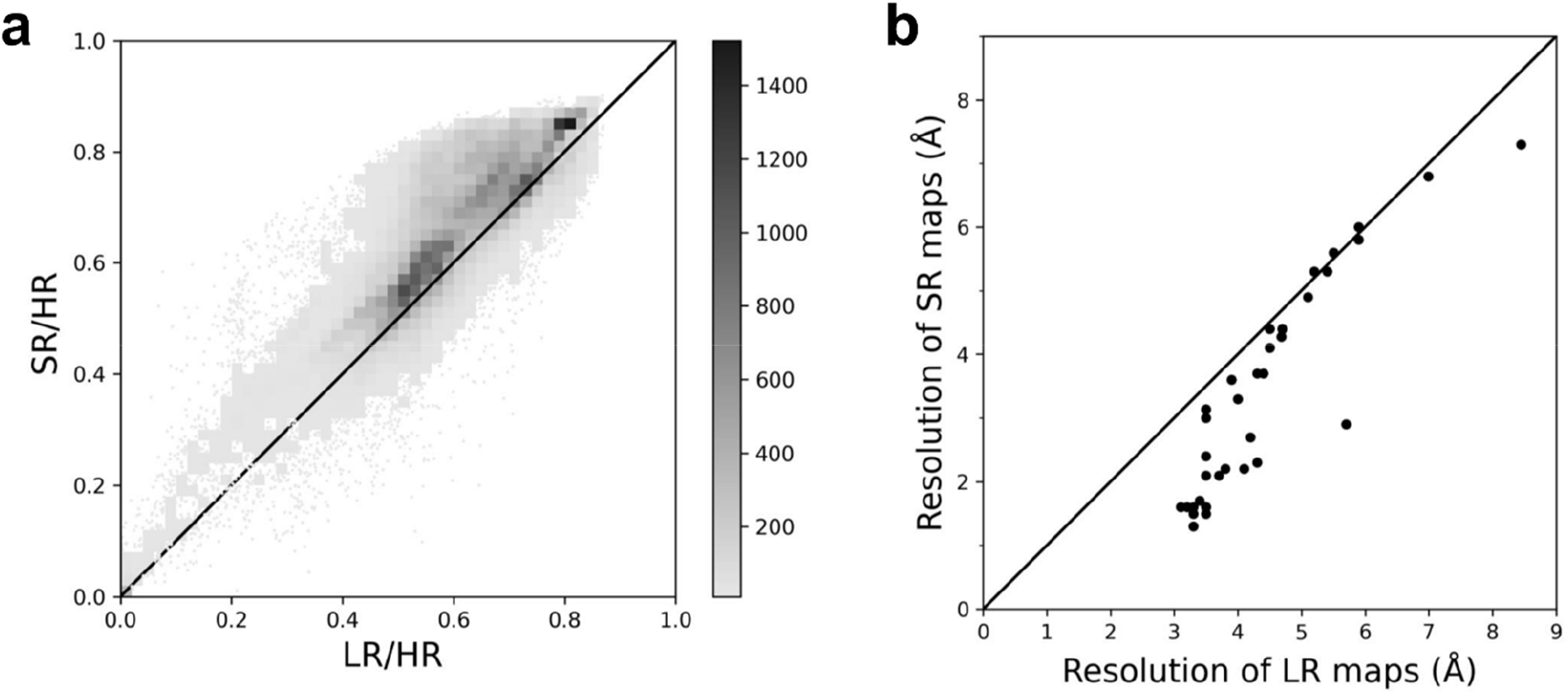
Local and global map resolution improvement by SuperEM. **a**, local resolution improvement. Cross-correlation of low-resolution & high-resolution (LR/HR) vs super-resolution & high-resolution (SR/HR) of local 91,197 cubes (36 test maps) shown. Gray scale bar indicates the number of points in each of the 50×50 bins in the plot. **b**, global resolution improvement. the estimated resolution using the “d_model” metric in the phenix.mtriage tool for each LR map and SR map.

**Figure 2a** shows that overall SuperEM improved the cross-correlation of cubes for most of the cases. An improvement in cross-correlation was observed for 84.3% (76895/91197) of the test cubes with an average improvement from 0.600 to 0.658 (9.67%). **Supplementary Figure 1** provides the same type of plots for the two map groups, maps that have an experimental resolution of 3.5 Å or better and simulated at 1.8 Å (**Supplementary Fig. 1a**) and those which have a resolution worse than 3.5 Å and simulated at 3 Å (**Supplementary Fig. 1b**). For both groups, almost the same fraction of the map cubes, 84.4% and 84.3% for the former and the latter groups, respectively, have improved cross-correlation to HR map cubes. Next, we examine the resolution improvement on a global map level. In **Figure 2b**, we generated full size SR maps by combining SR cubes. Average density values were assigned for overlapping regions of cubes. The map resolution of the experimental (LR) maps and SR maps where computed by the phenix.mtriage tool^31^ in the Phenix package^32^. It is apparent from **Figure 2b** that the resolution estimate of SR maps is clearly better over LR maps in 91.7% (33 out of 36) of the maps. The largest improvement was observed for map EMD-3672, which has an estimated resolution of 5.7 Å, which was improved by 2.8 Å to 2.9 Å. The exact resolution estimate values for all the test maps are provided in **Supplementary Table S1**.

### Examples of improvement by SuperEM

**Figure 3** shows six examples of EM maps before and after application of SuperEM. In each panel, three figures are shown: the native protein structure on the left, the initial LR EM map in the middle, and the SR EM map generated by SuperEM on the right. For all cases, the contour levels of LR and SR maps are visualized at the contour levels that give the similar density volumes in the two maps.

**Figure 3.**
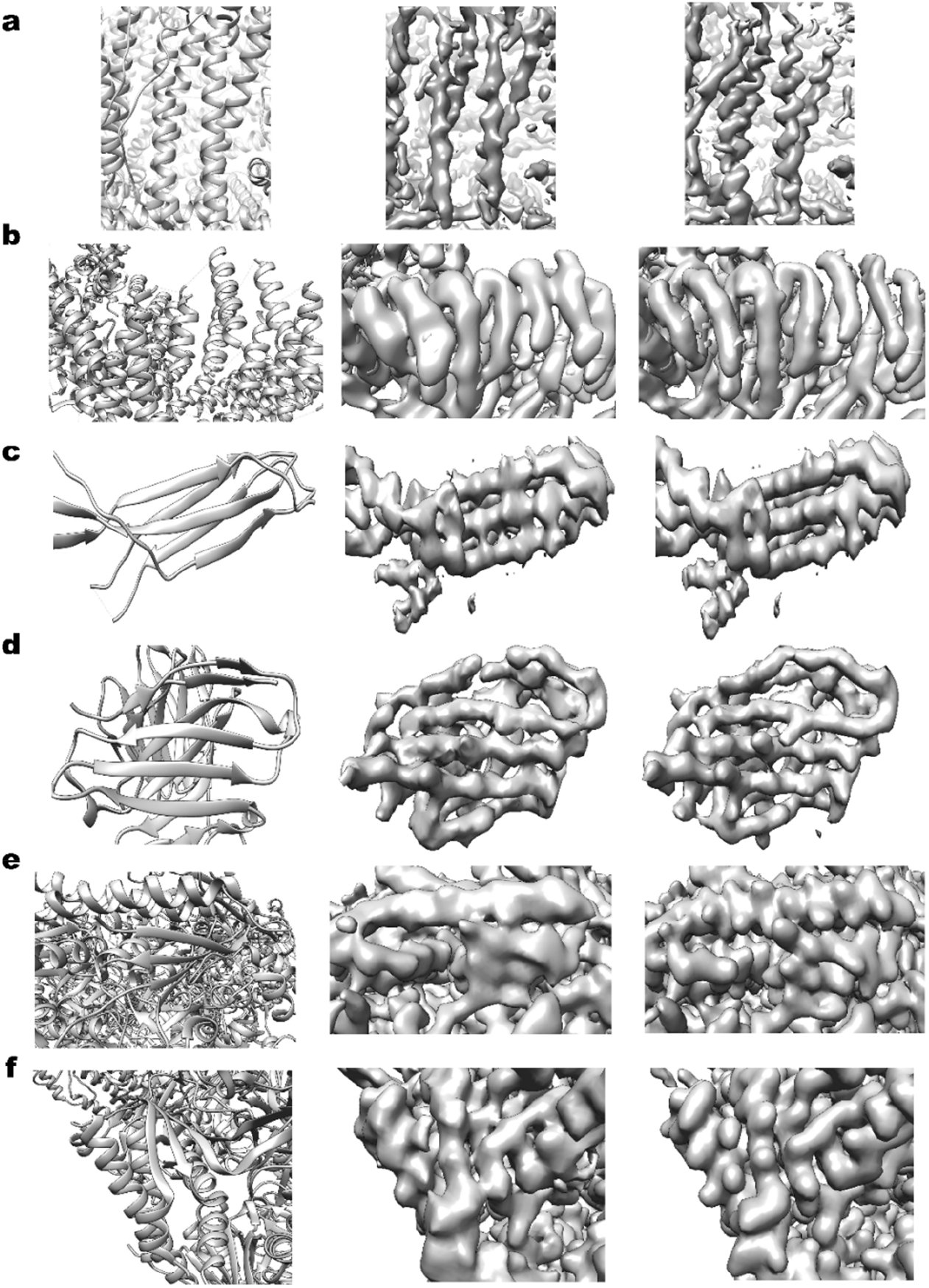
Examples of EM maps before and after applying SuperEM. In each panel, figure on the left are the native structure; middle, the experimental EM maps which here denoted as Low Resolution (LR) map; right, the map output from SuperEM denoted as Super-Resolution (SR) map. **a**, horse spleen apoferritin. EMD-2788, PDB ID: 4v1w. The resolution of the LR map was reported as 4.7 Å and measured by Phenix mtriage as 4.7 Å. The resolution of SR map measured by Phenix mtriage was 4.4 Å. The LR map visualized at a contour level of 0.270 and the SR map shown at a contour level of 0.320. These contour levels gave similar density volumes of the two maps, 50,839 Å^3^ for the LR map and 51,089 Å^3^ for the SR map. **b**, Tra1 subunit of SAGA complex. EMD-3790, PDB ID: 5oej. The resolution of the LR map: 5.7 Å/7.86 Å in EMDB/mtriage. The SR map: 6.99 Å. The contour levels used for visualization were 0.043 for the LR map, which gave a volume of 173,470 Å^3^ and 0.200 for the SR map, which gave a volume of 172,440 Å^3^. **c**, aerolysin, EMD-8188, PDB ID: 5jzw. The resolution of the LR map: 4.46 Å/4.5 Å in EMDB/mtriage. The SR map: 4.1 Å. The contour levels used for the LR map: 0.125 (16,271 Å^3^); the SR map: 0.220 (16,476 Å^3^). The density volume by using the contour level is shown in the parentheses. **d**, BG505 SOSIP.664 trimer in complex with HIV antibody 3BNC117. EMD-8644, PDB ID: 5v8m. The resolution of the LR map: 4.4 Å/4.4 Å in EMDB/mtriage. The SR map: 3.7 Å. The contour levels used for the LR map: 0.045 (108,480 Å^3^); the SR map: 0.244 (109,180 Å^3^). **e**, yeast Pol II transcription initiation complex. EMD-3378, PDB ID: 5fyw. The resolution of the LR map: 4.35 Å/4.55 Å in EMDB/mtriage. The SR map: 4.25 Å. The contour levels used for the LR map: 0.033 (235,110 Å^3^); the SR map: 0.180 (235,470 Å^3^). **f**, human TRPM4 channel in complex with calcium and decavanadate. EMD-8771, PDB ID: 5w5y. The resolution of the LR map: 3.8 Å/3.8 Å in EMDB/mtriage. The SR map: 2.2 Å. The contour levels used for the LR map: 0.040 (359,990 Å^3^); the SR map: 0.152 (359,830 Å^3^).

**Figure 3a** shows an α-class structure of horse spleen apoferretin^33^ (EMD-2788; PDB ID: 4v1w). The LR (middle) and SR (right) maps were shown at contour levels of 0.27 and 0.32, respectively, which give similar volumes of around 51,000 Å^3^. α-helices in the SR map have more distinct pitch than those in the LR map. Also, fragmented density regions originally in the LR map were connected following protein chains in the SR map. The next panel (**Figure 3b)** is another α helix-rich structure of Tra1 subunit of SAGA complex^34^ (EMD-3790, PDB ID: 5oej). In this example, SR map captures individual helices more distinctively than the LR map.

The next two panels illustrate the performance of SuperEM for maps of β-class proteins (**Figure 3c, 3d**). In both cases, β-strands are separated clearer in SR maps. Besides, in **Figure 3d**, a large gap in the density that correspond to a β-strand at the top of the map existed in the LR map but is connected in the SR map. The last two panels are from αβ-class proteins (**Figure 3e, 3f**). Compared to the LR maps, it is clear that large sidechains of amino acids are more visible in the SR maps.

### Structure modelling improvement by SuperEM

In the previous sections, we discussed differences made by SuperEM in density distributions of EM maps. Next, we examine how the map improvement translates practically to actual protein structure modeling. We used software for protein structure modeling, MAINMAST^3^ and Phenix^5^. To perform modeling, we first segmented a local density region that corresponds to the single subunit to model using the UCSF Chimera’s “zone” tool. The modeling software was then run on LR and SR maps independently. For MAINMAST, we ran the protocol up to the main-chain generation step. MAINMAST generates 15,000 models with different parameter combinations, from which one model was selected for evaluation based on the weighted edge distance along the trace^3^. Regarding Phenix, we used the map_to_model program, which takes an EM map, the target protein sequence, and a map resolution information to build a full atomic model. For the map resolution information to run map_to_model, we used the value reported in EMDB.

In **Figure 4**, we show the coverage of main-chain trace models generated by MAINMAST and models by Phenix in the two panels. Following previous works^3,5,35^, the coverage is defined as the fraction of Cα atoms in the native structure that are modelled within 3.0 Å in the model. Each panel compares the coverage of models generated from LR maps and SR maps of the 36 maps. For both methods, coverage improved for a majority of the cases by using SR maps. For MAINMAST, an improvement in coverage was observed for 24, while 22 cases showed improvement for Phenix and for 1 case the coverage stayed the same for both methods. 11 cases where the coverage decreased in MAINMAST models include maps of a low resolution, 6.0 Å (EMD-8581) and 5.8 Å (EMD-5779), but in general there is no clear correlation with the map resolution.

**Figure 4.**
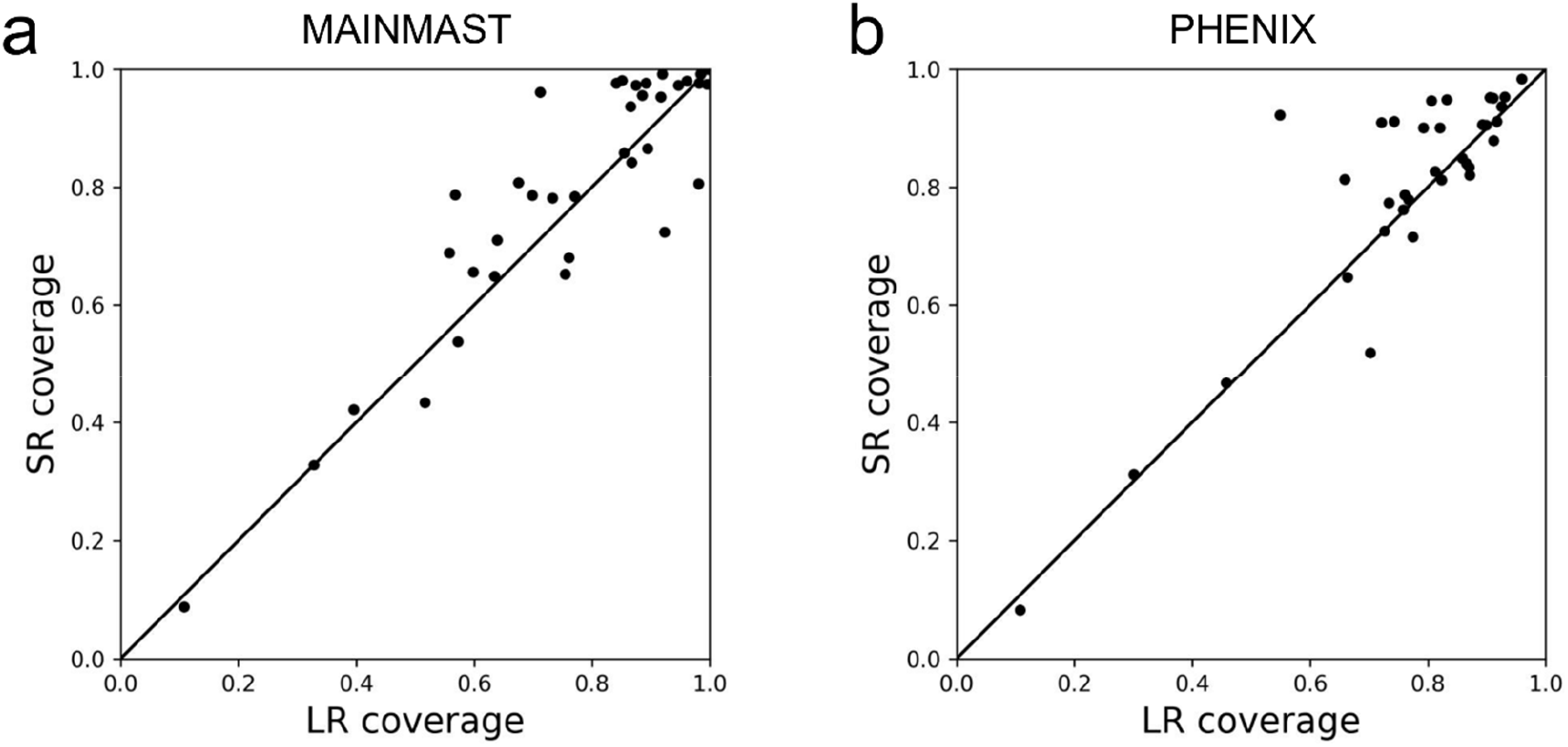
Change of the model coverage by two de novo modelling methods, MAINMAST and Phenix. Coverage within 3.0 Å are compared for models generated from LR and SR maps. **a**, models by MAINMAST. The number of cases that improved/tied/worsened by using SR maps were 24/1/11, respectively. **b**, models by Phenix. The number of cases that improved/tied/worsened by using SR maps were 22/1/11, respectively. There were two maps, EDM-2484 and EMD-8581, for which Phenix did not run because the protein sequence of the maps includes unknown amino acids (denoted as X).

**Figure 5** shows several examples of models generated for LR and SR maps by MAINMAST and Phenix. In practical scenario, modeling for maps at an intermediate resolution as shown here would certainly need many manual interventions such as template-based fitting, trying different parameters, and structure refinement. But here we show main-chain trace models from an automated run of the methods for the sake of the comparison of LR and SR maps. These models further need sequence mapping, full-atom building, and refinement with manual interventions. In each panel in **Figure 5**, the native structure in the original map (green), a model generated from the LR map (cyan), and a model from the SR map (magenta) are shorn from left to right. MAINMAST models are in the upper row and corresponding Phenix models are put in the lower row.

**Figure 5.**
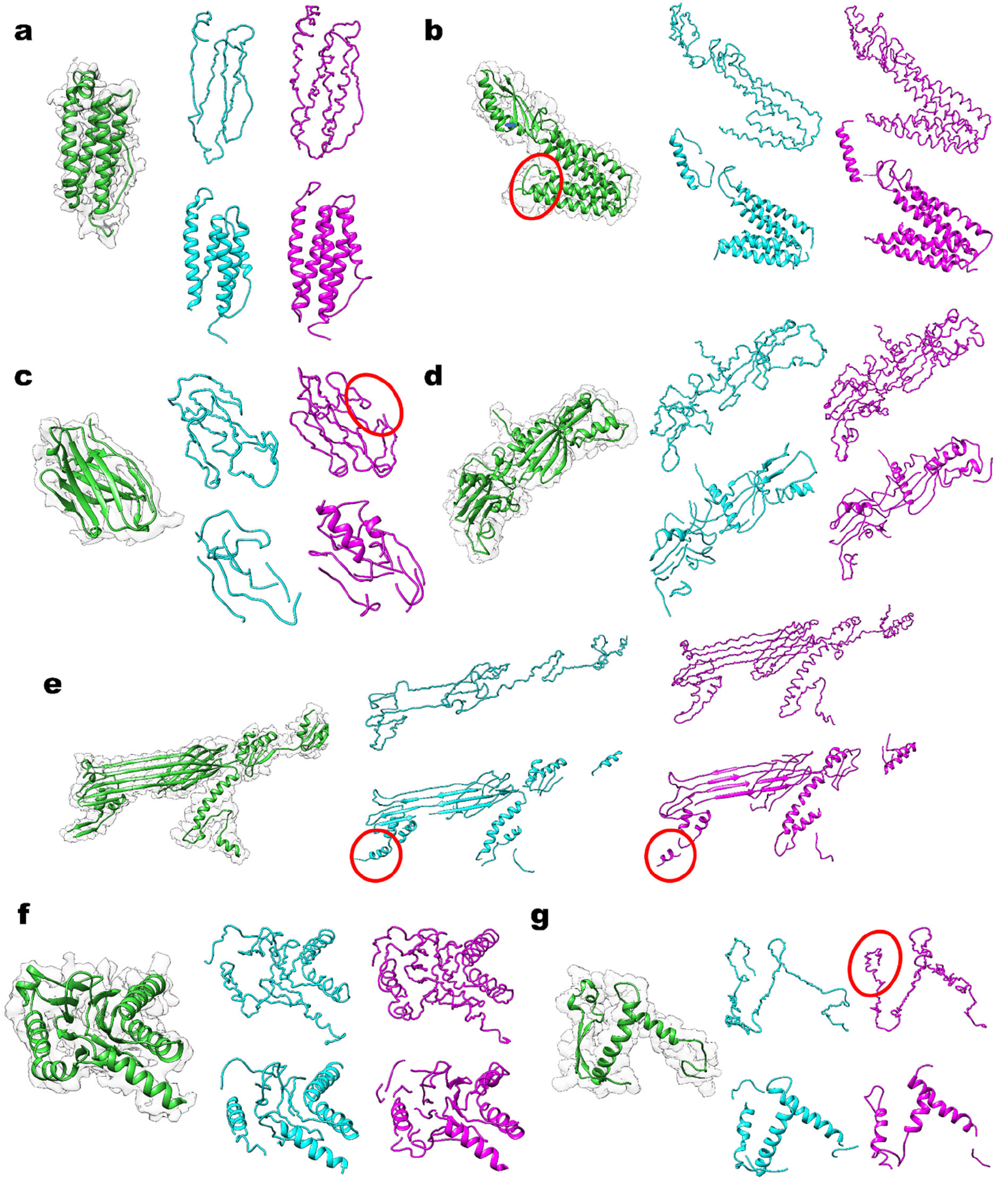
Examples of main-chain models by MAINMAST Phenix models from EM maps before and after SuperEM application. Each panel has five figures: Left most column, the experimental LR map superimposed with the corresponding protein structure in green. On the right, the upper row shows MAINMAST models the lower row shows Phenix models. Cyan, models from original LR maps; magenta, models from the SR maps. Local structures mentioned in the text are highlighted with red circles. **a**, A 174-residue chain B of horse spleen apoferritin (EMD-2788; PDB ID:4V1W). The map resolution of the LR map was 4.7/4.7 Å in EMDB/by Phenix mtriage. The resolution of the SR map: 4.4 Å by Phenix mtriage. The MAINMAST main-chain-trace model/the Phenix model generated from the LR map had a coverage within 3 Å of 0.841/0.806, respectively. From the SR map, the coverage of MAINMAST/Phenix models were 0.976/0.947, respectively. **b**, The zinc transporter Yipp (EMD-8728, PDB ID: 5VRF-B), 282 residue-long. The resolution of the LR map: 4.1 Å/4.1 Å. The SR map: 2.2 Å. Coverage of the MAINMAST/Phenix LR models: 0.894/0.823. The SR models: 0.865/0.812. **c**, heavy chain of NIH antibody 3BNC117 (EMD-8644, PDB ID: 5V8M-H), 111 residue-long. The resolution of the LR map: 4.4 Å/4.4 Å. The SR map: 3.7 Å. Coverage of the MAINMAST/Phenix LR models: 0.865/0.721. The SR models: 0.937/0.910. **d**, DNA-directed RNA polymerases I and III subunit RPAC1 (EMD-8771, PDB ID: 5W5Y-C), 306 residue-long. The resolution of the LR map: 3.8 Å/3.8 Å. The SR map: 2.2 Å. Coverage of MAINMAST/Phenix LR models: 0.712/0.899. The SR models: 0.961/0.905. **e**, the type II secretion system, chain D (EMD-6677, PDB ID: 5WQ9-D), 493 residue-long. The resolution of the LR map: 4.22 Å/4.2 Å. The SR map: 2.7 Å. Coverage of the MAINMAST/Phenix LR models: 0.568/0.734. The SR models: 0.787/0.773. **f**, bacterial proteasome subunit α (EMD-4128, PDB ID: 5LZP-Y), 213 residue-long. The resolution of the LR map: 3.5 Å/3.5 Å. The SR map: 1.5 Å. Coverage of the MAINMAST/Phenix LR models: 0.851/0.930. The SR models: 0.981/0.953. **g**, Ebola surface glycoprotein, GP2 (EMD-8242, PDB ID: 5KEN-M), 113 residue-long. The resolution of the LR map: 4.3 Å/5.5 Å. The SR map: 5.6 Å. Coverage of MAINMAST/Phenix LR models: 0.761/0.726. The SR models: 0.681/0.726.

The first example (**Figure 5a**) is modeling for a 174-residue long α class protein. The original resolution was 4.7Å, which was improved to 4.4 Å by SuperEM. Phenix built very good models for this case (the lower row in the panel). The SR map made it possible for Phenix to build a structure with the correct topology, which has four long α helices placed at correct positions with minor errors of missing a short helix and an overprediction of another helix. The Phenix model from the LR map has a break in the chain and made a helix shorter, but they were fixed when the SR map was used. MAINMAST traced the main-chain almost correctly, with one wrong loop connection at the bottom of the structures. By using the SR map, the local geometry of helices substantially improved in the MAINMAST models. Apparently, the quality of helices is low in MAINMAST models but it is to be corrected in the subsequent structure refinement step in the MAINMAST protocol^3^. The second example (**Figure 5b**) has essentially the same observation. This protein has two domains, an α-helix bundle (on the right side of the figure) and a small α/β domain with a β-sheet and an α helix. By using the SR map, MAINMAST improved the helix geometry and improved the coverage in the α/β domain. The two Phenix models from the LR and SR maps have some differences, but the overall quality were similar. All the models missed a loop connection indicated by a circle in the figure because the local map density of the position is relatively low.

The next three panels, c, d, e are examples of modeling of β-sheets. **Figure 6c** is models for an antibody heavy chain. In this case, the MAINMAST results substantially improved by using the SR map, where the main-chain was almost perfectly traced except for a missing connection on the right side of the structure (circled). On the other hand, the two Phenix models were fragmented and moreover, the SR model has two helices that were misassigned. The next example is (**Figure 6d**), a structure of a complexed topology relative to the other structures shown in **Figure 6**. MAINMAST showed an excellent performance for this example, where the main-chain was perfectly traced in the SR map with small non-critical deviations, achieving a high coverage of 0.961. In both MAINMAST and Phenix, models from the SR maps showed clear improved over those from LR maps. **Figure 6e** is a subunit from Type II secretion system, which has a characteristic large β-sheet associated with another smaller β-sheet behind the large one. Modeling by MAINMAST largely improved with the SR map, which built the correct topology for the domain with the two β-sheets and a long helix in the middle of the structure. Phenix also corrected the topology of the small β-sheet with the SR map and has a larger coverage in the two domains on the right. A small helix was wrongly assigned at the left bottom of the structure (red circle) by Phenix, which still existed in the SR model.

**Figure 6f** is an example of α/β proteins, which have a two-layer β sheets in the middle and five surrounding α helices. With MAINMAST, the entire topology is almost perfectly traced with the SR map whereas a β-sheet on the left is non-existent in the LR model. These structural enhancements resulted in a coverage increase of from 0.851 (the LR model) to 0.981 (the SR model). The models by Phenix are fragmented but the SR model captures the core part of the topology correctly.

The last panel (**Figure 6g**) is an example where the modeling did not improve with the SR map. This protein has a two-stranded β-sheet on the left connected with two helices on the right. For this map, SuperEM made the map resolution worse, from 5.5 Å originally to 5.6 Å. Partly for this reason, modeling results by both MAINMAST and Phenix saw essentially no improvement. Neither method could trace the β-sheet correctly, placing an α helix in the middle of the two strands (Phenix) or intersected two strands (MAINMAST). MAINMAST dropped the coverage from 0.761 to 0.681 by missing a longer loop region on the left top of the figure.

### Comparison with a related method

Lastly, we compared SuperEM with DeepEMhancer^11^, another similar deep-learning-based method for sharpening of EM maps. DeepEMhancer is different from SuperEM in that it aims to produce local sharpening effects of the LocScale^8^ algorithm using a 3D U-Net^36^, a different neural network architecture. Since the goals of the two methods are similar but not identical, this comparison is solely to characterize the performance of SuperEM.

The performance comparison is provided in **Supplementary Table S2**. We tested the two methods on 17 EM maps, which have half-maps provided at EMDB, because the preferred input for DeepEMhancer is half-maps as their deep-learned model was trained with half maps. There are six maps with half-map data in the 36 maps in the dataset we used. To increase the dataset size, we added 11 maps that were deposited recently to EMDB (after May 2020) with half-maps made available. SuperEM showed better resolution than DeepEMhancer for 14 out of 17 cases. The average resolution improvement by SuperEM was 1.35 Å while that for DeepEMhancer was 0.57 Å.

## Discussion

In this work, we developed SuperEM, a novel 3D deep learning-based framework to produce super-resolution cryo-EM maps. The improvement of resolution was possible for almost all the maps examined, and the resolution improvements were translated into more accurate protein structure modeling by two modeling methods. The current framework can be easily extended to maps at higher or lower resolutions by training the network with an appropriate dataset. Thus, together with other experimental and map building techniques for enhancing the density map quality, SuperEM will be a valuable asset for achieving better quality structures.

## Acknowledgments

The authors would like to acknowledge Thomas Klose for helpful discussions. The authors are grateful to Charles Christoffer for his help in preparing the manuscript and Xiao Wang for his feedback for the codes released on Github. This work was partly supported by the National Institutes of Health (R01GM123055, R01GM133840), the National Science Foundation (DMS1614777, CMMI1825941, MCB1925643, DBI2003635), and the Purdue Institute of Drug Discovery.

## Author Contributions

D.K. conceived the study. S.R.M.V.S. developed the algorithm and coded SuperEM. The datasets were selected by S.R.M.V.S. and G.T. The experiments were designed by S.R.M.V.S. and D.K. and were carried out by S.R.M.V.S. S.R.M.V.S., G.T., and D.K. analyzed the results. The manuscript was drafted by S.R.M.V.S. D.K. administrated the project and wrote the manuscript.

## Code availability

The SuperEM program is freely available for academic use through https://github.com/kiharalab/SuperEM and https://kiharalab.org/emsuites/superem.php.

## Data availability

The data that support the findings of this study are available from the corresponding author upon request.

## Methods

### HDataset Generation

SuperEM requires a dataset of pairs of cubes of EM density maps, low-resolution and high-resolution, for training. We used cubes extracted from experimental EM maps as low-resolution data and their corresponding simulated EM maps from associated atomic-detailed structures from PDB as high-resolution data.

The data generation is similar to the procedure of the experimental map data generation performed for Emap2sec^37^, another deep-learning method for cryo-EM map data processing. For the low-resolution dataset, we obtained experimental EM maps from EMDB as follows: 788 EM maps with resolution between 3 Å–6 Å dated up to December 2018 were first selected. These EM maps also had an associated atomic structure available in PDB so that we can generate the corresponding simulated EM map. We further short-listed 377 maps, which have the cross-correlation between the experimental map and the map simulated from the corresponding PDB structure over 0.65. We then computed the sequence identity between underlined protein chains in every pairs of EM maps, and a map was removed from the dataset if its underlined protein had over 25% identity to a protein of another map in the dataset. If a map has multiple protein chains, the map was removed if at least one of the chains have over 25% identity to any chains in another map. This procedure finally remained 122 experimental EM maps. We unified the grid sizes of the experimental maps to 1.0 Å by trilinear interpolation of the electron density in the maps. High-resolution simulated EM maps were generated from the PDB structures of the above 122 experimental maps using pdb2vol program from SITUS package^38^. For experimental maps with resolution ranging between 3.5 Å and 6.0 Å, the corresponding high-resolution simulated map was generated at 3.0 Å. For experimental maps between 3 Å and 3.5 Å, a high-resolution was simulated at 1.8 Å.

In both simulated and experimental maps, electron density values were normalized from 0.0 to 1.0 by subtracting the minimum density value in the map and then divide by the density difference between the maximum and the minimum density values. If there are negative density values, they were set to 0.0 before the normalization. Each of the EM maps were then converted to voxels by a sliding cube of size 25 Å *25 Å *25 Å moved through the maps with a stride size of 4 Å. This procedure generated a total of 292,749 valid low-resolution and high-resolution pair of cubes. A cube pair is considered valid if neither of the cube contains all-zero density values. Out of 122 EM map pairs generated, 86 pairs amounting to 201,552 cube pairs were used for training and 36 pairs amounting to 91,197 cube pairs were used for testing. Among the 36 maps in the testing set, 12 of them have a resolution better than 3.5 Å and 24 maps had a resolution worse than 3.5 Å.

### Training procedure of SuperEM

The training cube pairs described above were used to train the GAN network of SuperEM. Among the cube pairs, the cube from the experimental map, which we here-by refer to as a low-resolution (LR) cube, is the input to the Generator network of GAN. The output cube of generator network and the corresponding cube from simulated map or the high-resolution (HR) cube are then fed to the discriminator network.

In GANs, the generator and discriminator are trained together with a minimax game-style objective function given by Equation 1.

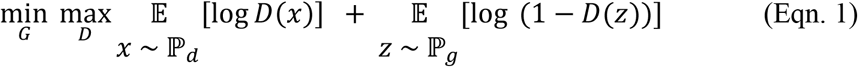

where G and D are parameters of generator and discriminator networks of GAN, ℙ_*d*_ is HR cube distribution, ℙ_*g*_ is the generated cube distribution. The generator receives an LR cube as input and generates a super-resolution (SR) cube. The discriminator then classifies the SR cube and HR cube. The minimax objective ensures that the generator generates superior quality SR cubes so that the generator can fool the discriminator into classifying them as HR cubes. The objective function for GAN is formulated, in Equation 2, as a linear combination of content loss and adversarial loss which are given in Equations 3 and 4.

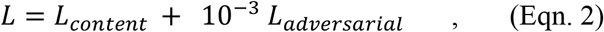

Where

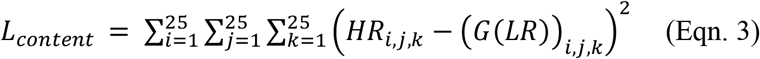

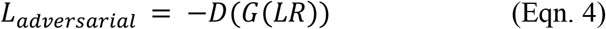

where HR corresponds to the high-resolution cube, and LR corresponds to the low-resolution cube which makes G(LR) the super-resolution cube and D(G(LR)) is the discriminator probability output for the SR cube ranging from 0 to 1. Negative of D(G(LR)) is optimized to fool the discriminator to think that SR cube is as good as the HR cube. To summarize, the content loss is defined by the Mean Squared Error (MSE) between the SR cube and the HR cube. The adversarial loss is given as the negative softmax probabilities of the discriminator predictions.

We used a learning rate of 0.001 for both the generator and the discriminator. We used a batch size of 16 and train the GAN network for 100 epochs using the Adam Optimizer for updating the weights of both the generator and the discriminator.

## Supplementary Information for

**Supplementary Figure 1.**
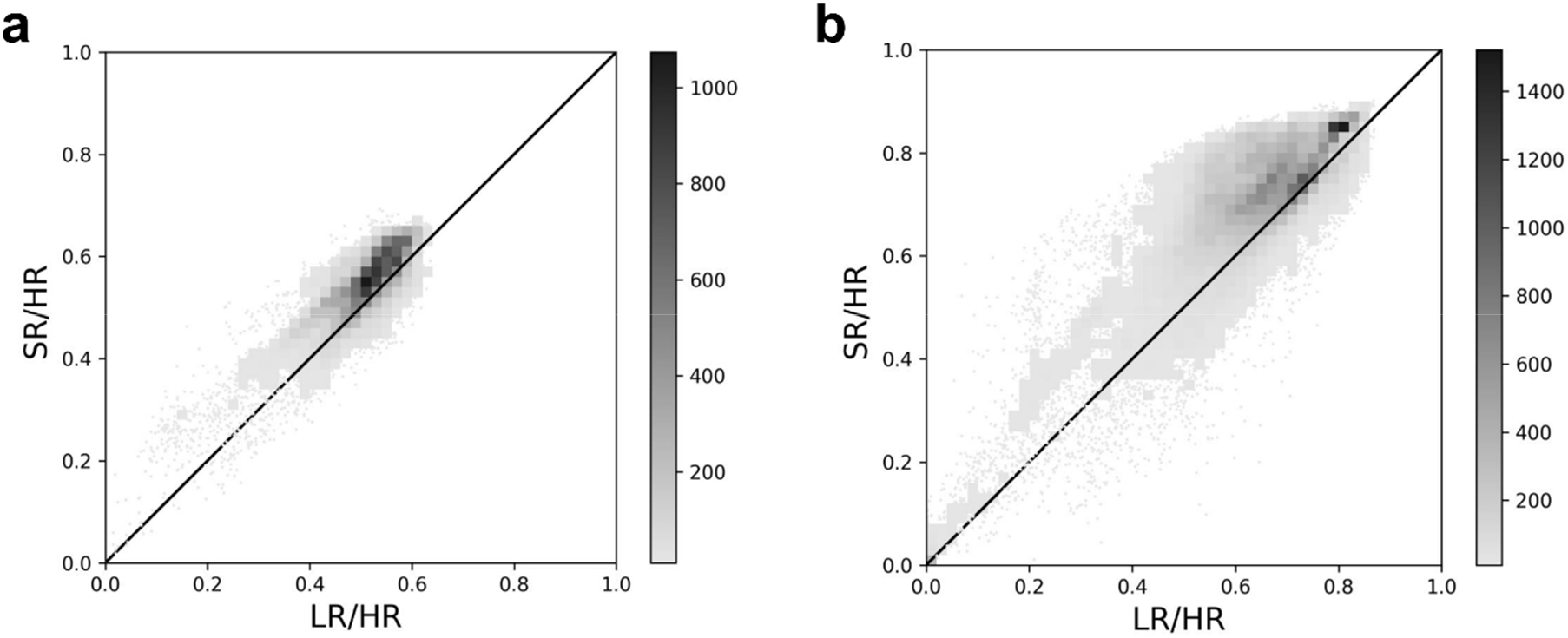
Local resolution improvement of cryo-EM maps by SuperEM at two different simulated resolutions. Gray scale bar indicates the number of data points in each bin of the histogram consisting of 50×50 bins. White indicates a bin containing less than 10 data points. **a**, Cross-correlation of low-resolution & high-resolution (LR/HR) vs. super-resolution & high-resolution (SR/HR) of 25,565 map cubes for 12 EM maps with an experimental map resolution better or equal to 3.5 Å. The corresponding simulated maps were computed at 1.8 Å. 84.4% (23257/27565) of the cubes had an improved cross-correlation with an average improvement from 0.508 to 0.544 (7.09%). **b**, Cross-correlation of LR/HR vs SR/HR of 63,632 cubes for 24 EM maps with experimental map resolution worse than 3.5 Å. The corresponding simulated maps were computed at 3.0 Å. 84.3% (53638/63632) of the test cubes improved with an average improvement from 0.640 to 0.707 (9.78%).

**Supplementary Table 1. (provided in a separate Excel file). Detailed results of the resolution estimate value for 36 test maps**. Columns in the table are: Map EMID, corresponding PDB code, map resolution in Å taken from EMDB, Phenix mtriage resolution estimates in Å for each of low-resolution (LR) i.e. experimental EM map, high-resolution (HR) i.e. simulated EM map and Super-Resolution (SR) EM map are provided. Last column shows the improvement in resolution of SR map compared to LR map.

**Supplementary Table 2.** Comparison of resolution estimates between EM maps generated by SuperEM and DeepEMhancer.

| EMDB-ID | Original | SuperEM | DeepEMhancer |
| --- | --- | --- | --- |
| 3663* | 5.16/5.90 | 5.80 | **5.50** |
| 4078* | 5.10/5.10 | 4.90 | **4.20** |
| 5623* | 3.30/3.30 | **1.50** | 2.90 |
| 6479* | 3.50/3.50 | **1.60** | 3.30 |
| 6714* | 3.00/3.10 | **1.60** | 2.90 |
| 8624* | 3.40/3.50 | **2.10** | 3.00 |
| 10649 | 4.30/4.30 | **2.60** | 4.00 |
| 10094 | 3.34/3.40 | **1.40** | 2.10 |
| 0975 | 3.10/3.50 | **1.80** | 3.00 |
| 0882 | 3.30/3.40 | **1.40** | 3.00 |
| 0920 | 3.40/3.70 | **1.80** | 3.10 |
| 0510 | 3.63/3.50 | **3.10** | 3.40 |
| 0981 | 2.84/3.20 | **1.40** | 2.00 |
| 10088 | 4.20/4.20 | **2.30** | 3.90 |
| 10350 | 3.70/4.00 | 3.70 | **3.10** |
| 10704 | 4.08/4.10 | **3.30** | 3.90 |
| 10836 | 3.30/3.40 | **1.80** | 2.10 |
| Avg. Gain | - | 1.35 | 0.57 |

Resolution estimate (Å) computed with phenix.mtriage tool (http://www.phenix-online.org/documentation/reference/mtriage.html) for maps generated by SuperEM and by DeepEMhancer are shown. Original shows the map resolution of the experimental maps. Two values are the one stated in EMDB/measured by phenix.mtriage. Comparing the resolutions by SuperEM and DeepEMhancer, the smaller value was highlighted in bold.
The six EMDB-IDs marked with \* are included in our benchmark set. The rest of the targets used here selected from recent deposits (after May 2020) to EMDB. All these targets have half-map data available at EMDB. Input to DeepEMhancer for a target map was half-map because it was trained on half-map and that is stated as the preferred input, while the input for SuperEM is the original deposited map to EMDB, as was used for the rest of the current study.
Avg. gain shows the average improvement of the resolution relative to the mtriage-measured resolution of the original map.

**Table 1.**
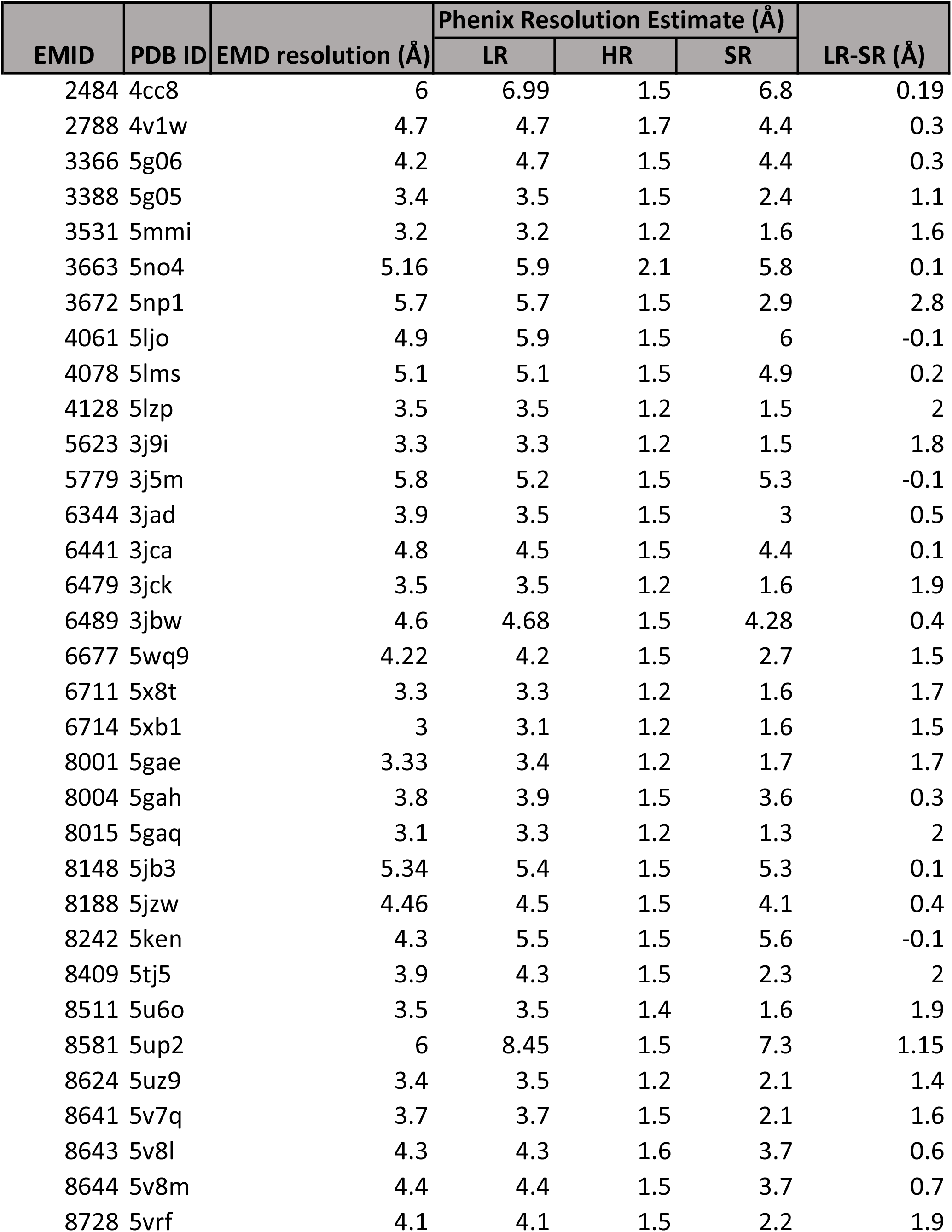
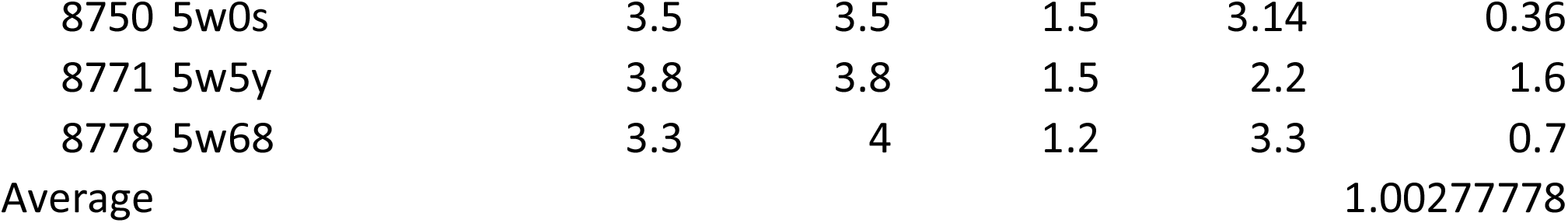
Detailed results of the resolution estimate value for the 36 maps.

